# An integrated workflow for long-term fiber photometry analysis

**DOI:** 10.64898/2026.04.21.719944

**Authors:** Farina Pourmir, Jordan N. Cook, Samantha O. Sweck, Jeff R. Jones

## Abstract

Long-term fiber photometry enables measurement of neural dynamics across hours to days, but these recordings create analytical and reproducibility challenges that are not well addressed by tools developed for short, stimulus-locked experiments. Here we present a software environment for long-term photometry analysis organized around a structured, revisitable workflow for run execution, inspection, and post-run refinement. The software separates correction retuning from downstream event reanalysis, allowing both signal correction and event-analysis settings to be revised after the initial run. We show that correction choice can substantially change the corrected signal itself and that post-run reanalysis can revise event-detection outcomes. The software also preserves tonic and phasic outputs and supports inspection of the same recording at both multiday and session-level scales. Together, these capabilities provide a practical workflow for more interpretable, revisitable, and reproducible analysis of long-term photometry recordings.

## INTRODUCTION

Genetically-encoded fluorescent indicators have expanded the temporal reach of systems neuroscience by making it possible to monitor neural activity across hours, days, and in some cases, weeks [1]. Many fundamental physiological and behavioral processes unfold over these longer timescales, shaped by internal state, environmental context, and recurring biological rhythms rather than by brief stimulus-evoked events [2– 7]. Circadian systems make the importance of these longer timescales especially clear, because their underlying neural and molecular dynamics unfold across the full 24 h cycle [8–13]. More broadly, the same challenge extends to any circuit whose activity changes slowly or recurrently over time. Long-term optical recording therefore offers direct access to aspects of neural function that were previously difficult to study.

Analytical assumptions that work well in short, stimulus-locked recordings often break down in long-term photometry datasets. Over these longer timescales, fluorescence reflects a mixture of true neural dynamics, drift, photobleaching, motion-related artifacts, and other slow biological or technical influences that complicate interpretation [14]. As a result, similar datasets can yield divergent conclusions when they are analyzed under different assumptions or through poorly standardized workflows. Long-term photometry therefore poses a problem of interpretation and reproducibility at the level of the analytical workflow itself.

Prior work from our lab used circadian photometry as a case study to argue that long-term optical monitoring requires analytical strategies tailored to extended recordings [1]. That paper emphasized reproducibility and standardization and introduced tonic and phasic decomposition to distinguish slow and transient components of long-duration fluorescence signals [15]. It further argued that common analysis practices often collapse meaningful slow structure into baseline drift or otherwise blur the distinct information carried by those signal components. Together, these ideas defined an analytical framework for reproducible long-term photometry analysis.

Long-term analysis places practical demands on software that go beyond the scope of most short-timescale workflows. Those demands include scalability, modularity, standardization, automated handling of large numbers of sessions, robust modeling of control signals, and workflow practices that preserve tonic and phasic structure. Existing open-source tools support important parts of fiber photometry analysis, especially for shorter experiments, but they do not fully solve the long-term workflow problem posed by extended recordings [16– 19]. Addressing this problem requires a practical, integrated analysis environment that makes reproducible long-term analysis feasible in routine laboratory use.

The software presented here was designed as an integrated analysis environment for long-term photometry, with particular emphasis on the analytical and workflow demands of extended recordings. It preserves the distinction between tonic and phasic signal structure, treats correction as a revisitable analytical stage rather than as a hidden preprocessing step, and separates correction retuning from downstream event reanalysis. More broadly, it provides a practical implementation of long-term photometry analysis principles within a reproducible software workflow.

## RESULTS

### A structured and revisitable workflow for long-term photometry analysis

We organized the software as a structured workflow for long-term photometry analysis, in which recordings are configured, validated, analyzed and then revisited through subsequent analysis iterations (**Figs. 1**,**2**). Within this workflow, completed runs can be reopened, reviewed, and revised rather than being treated as final after the initial run (**Fig. 1**). At a high level, this revisitable structure includes two distinct post-run branches, correction retuning and downstream event reanalysis, which are separated within the workflow and allow tuned outputs and settings to be applied in later runs.

**Figure 1.**
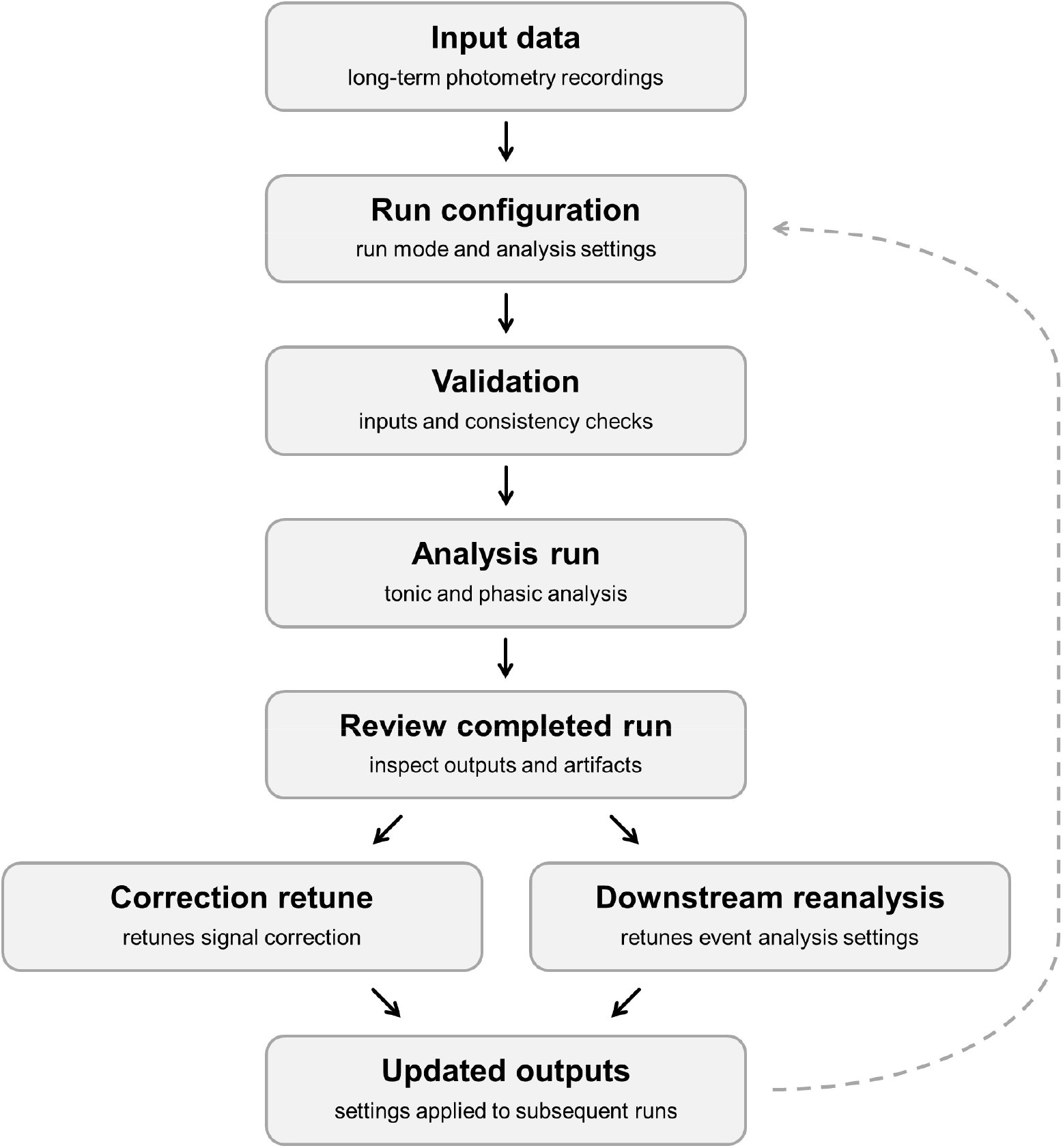
Workflow of the long-term photometry analysis environment. Long-term photometry data are configured, validated, and analyzed within a structured workflow that supports iterative refinement after completion of the initial run. Completed analyses can be reviewed and revisited through two distinct post-run branches: correction retuning, which modifies signal correction, and downstream reanalysis, which revisits event-analysis settings. Updated output and tuned settings can be applied to subsequent runs, enabling repeated refinement across analysis iterations.

This workflow is exposed directly through the graphical interface (**Fig. 2**). The setup environment is divided into standard and advanced branches, with standard windows for routine run setup and advanced windows for lower-level analysis settings. The standard branch contains the primary windows needed to initiate a run, including run configuration, ROI selection, and plotting and timeline-anchor settings (**Fig. 2a**). The advanced branch exposes controls for isosbestic correction, preprocessing, feature detection, and custom configuration loading (**Fig. 2b**).

**Figure 2.**
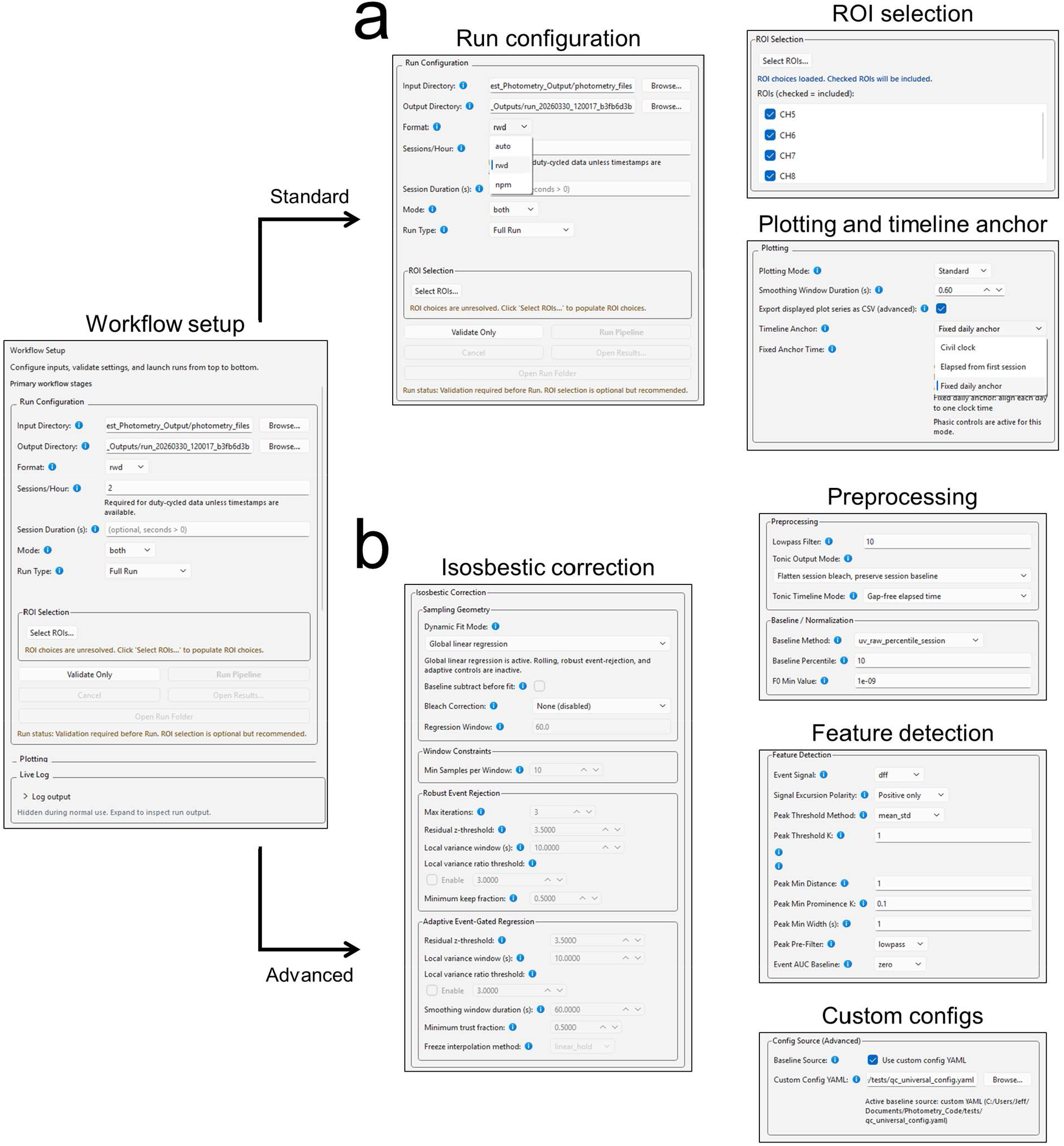
Standard and advanced setup windows in the long-term photometry analysis GUI. The workflow setup interface is organized into standard and advanced branches. **a)** The standard branch contains the primary windows used for routine run setup, including run configuration, ROI selection, and plotting and timeline-anchor settings. **b)** The advanced branch contains analysis parameterization windows for isosbestic correction, preprocessing, feature detection, and custom configuration loading.

### Execution progress and completed-run inspection within the GUI

The GUI reports progress through the major stages of the analysis workflow, including validation, tonic analysis, phasic analysis, and plotting (**Fig. 3a**). Progress is displayed directly within the interface as the run advances through these stages.

**Figure 3.**
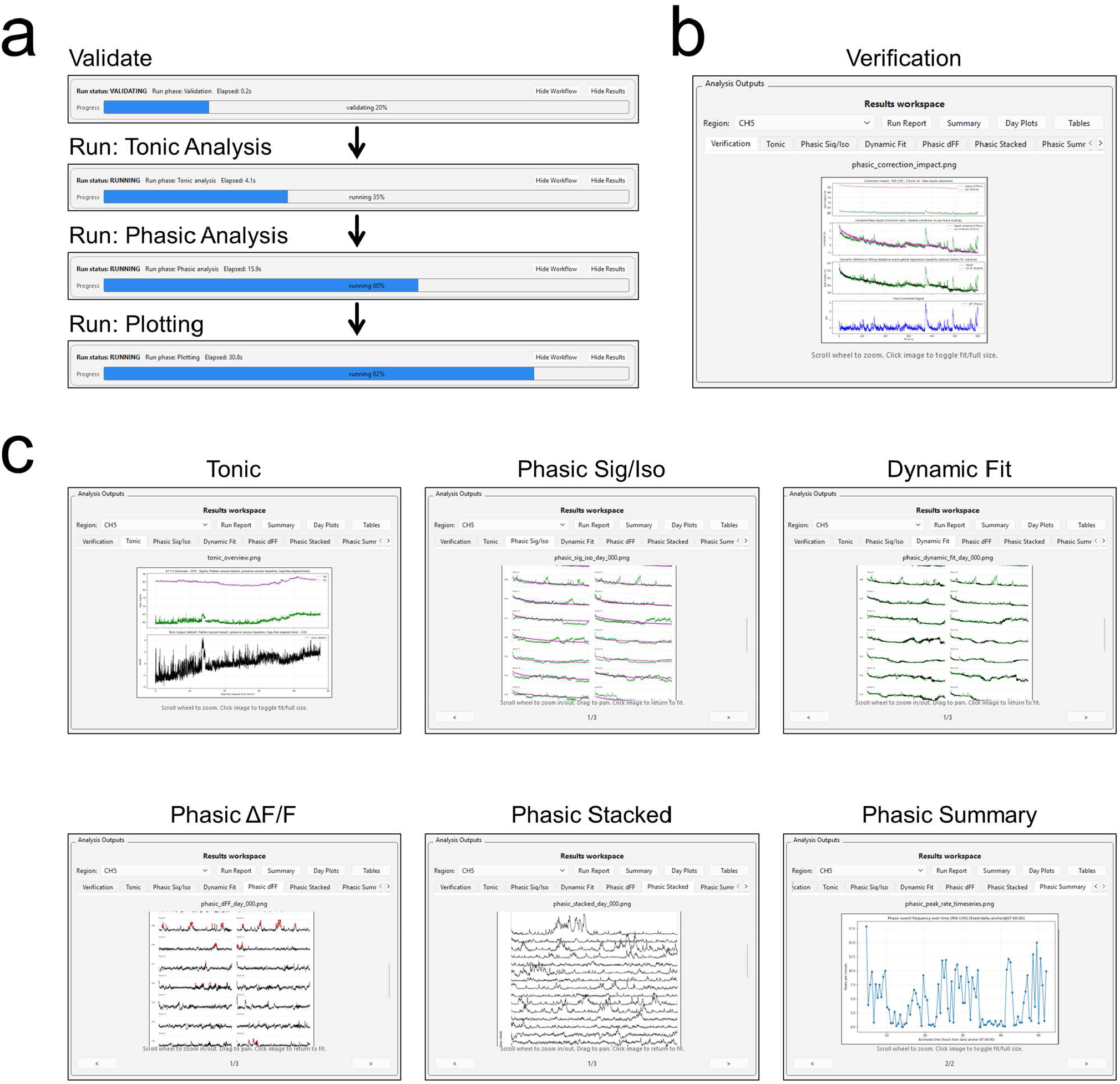
Execution progress and analysis output windows in the long-term photometry GUI. **a)** Representative progress-bar views from the analysis workflow, showing progression through validation and the main execution stages of tonic analysis, phasic analysis, and plotting. **b)** Analysis Output window after completion of a run, shown on the verification tab, which summarizes correction quality for review. **c)** Additional Analysis Output window tabs showing the main categories of post-run inspection available within the GUI, including tonic outputs, phasic signal and isosbestic views, dynamic-fit inspection, corrected phasic ΔF/F output, stacked phasic traces, and phasic summary views.

After completion of a run, outputs are available directly within an Analysis Output window for immediate inspection (**Fig. 3b,c**). The Verification tab provides a dedicated view for inspection of correction quality, while additional tabs expose the main categories of post-run outputs, including tonic outputs, signal and isosbestic views, dynamic fit inspection, corrected phasic ΔF/F, stacked phasic traces, and phasic summary outputs.

Completed runs can therefore be inspected within the same environment used to configure and execute the analysis.

### A dedicated post-run workflow for correction retuning

Because long-term recordings can be highly sensitive to correction assumptions, the software allows baseline and signal correction to be recomputed after completion of the initial run (**Fig. 4a**). This correction-retune workflow recomputes baseline and signal correction for a selected ROI, allowing correction to be revisited as an explicit analytical step after the initial run.

**Figure 4.**
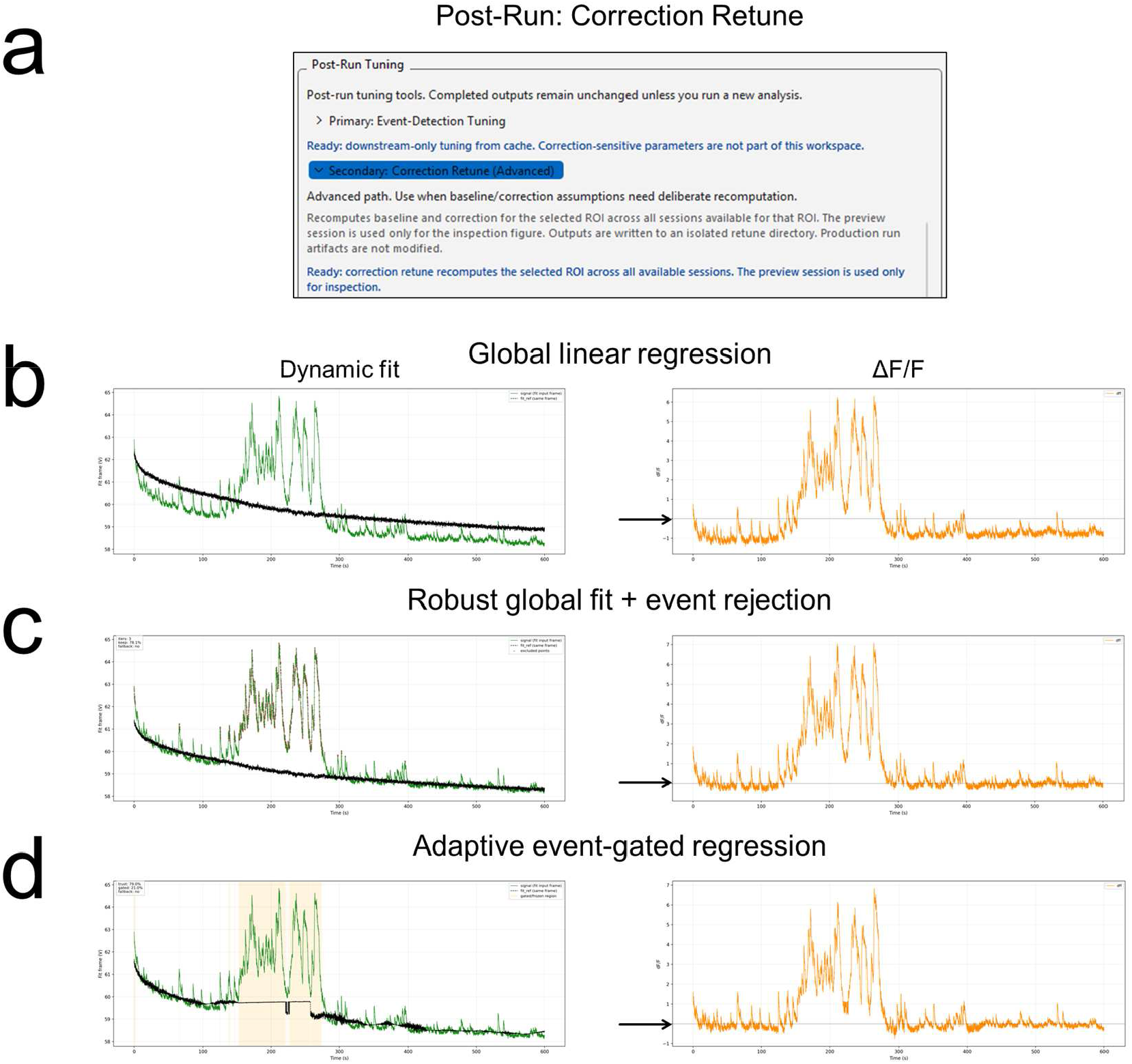
Post-run correction retuning and representative isosbestic correction outcomes. **a)** Post-run tuning window for correction retune, which recomputes baseline and signal correction for a selected ROI after completion of the initial run. **b)** Representative comparison of three isosbestic correction strategies applied to the same session. For each method, the left panel shows the signal trace and fitted isosbestic, and the right panel shows the resulting ΔF/F trace. Global linear regression mis-centers the corrected trace below the zero baseline (arrow), whereas robust global fitting with event rejection and adaptive event-gated regression both preserve an appropriate baseline near zero by preventing the large event from dominating the fit. The adaptive method also adjusts the fitted isosbestic locally across the session.

The need for this workflow is illustrated by comparing three isosbestic correction strategies applied to the same session (**Fig. 4b**). Under global linear regression, the fitted isosbestic is pulled upward by the large event, causing the corrected ΔF/F trace to become mis-centered below zero through most of the session. In contrast, both robust global fitting with event rejection and adaptive event-gated regression preserve a baseline near zero by preventing the large event from dominating the fit. In this example, the adaptive method also adjusts the fitted isosbestic locally across the session. These comparisons show that correction choice can substantially change the corrected signal itself, making correction retuning a necessary post-run step in the workflow.

### A dedicated post-run workflow for downstream event reanalysis

The software also supports post-run revision of downstream event analysis settings from cached phasic traces after completion of the initial run (**Fig. 5a**). In this branch, event analysis settings can be revisited without recomputing upstream signal correction. This workflow is illustrated on the same ΔF/F trace before and after downstream event detection retuning (**Fig. 5b**). Under the initial settings, excessive peaks are detected across the session. After retuning, a more appropriate set of peaks are detected across the same trace.

**Figure 5.**
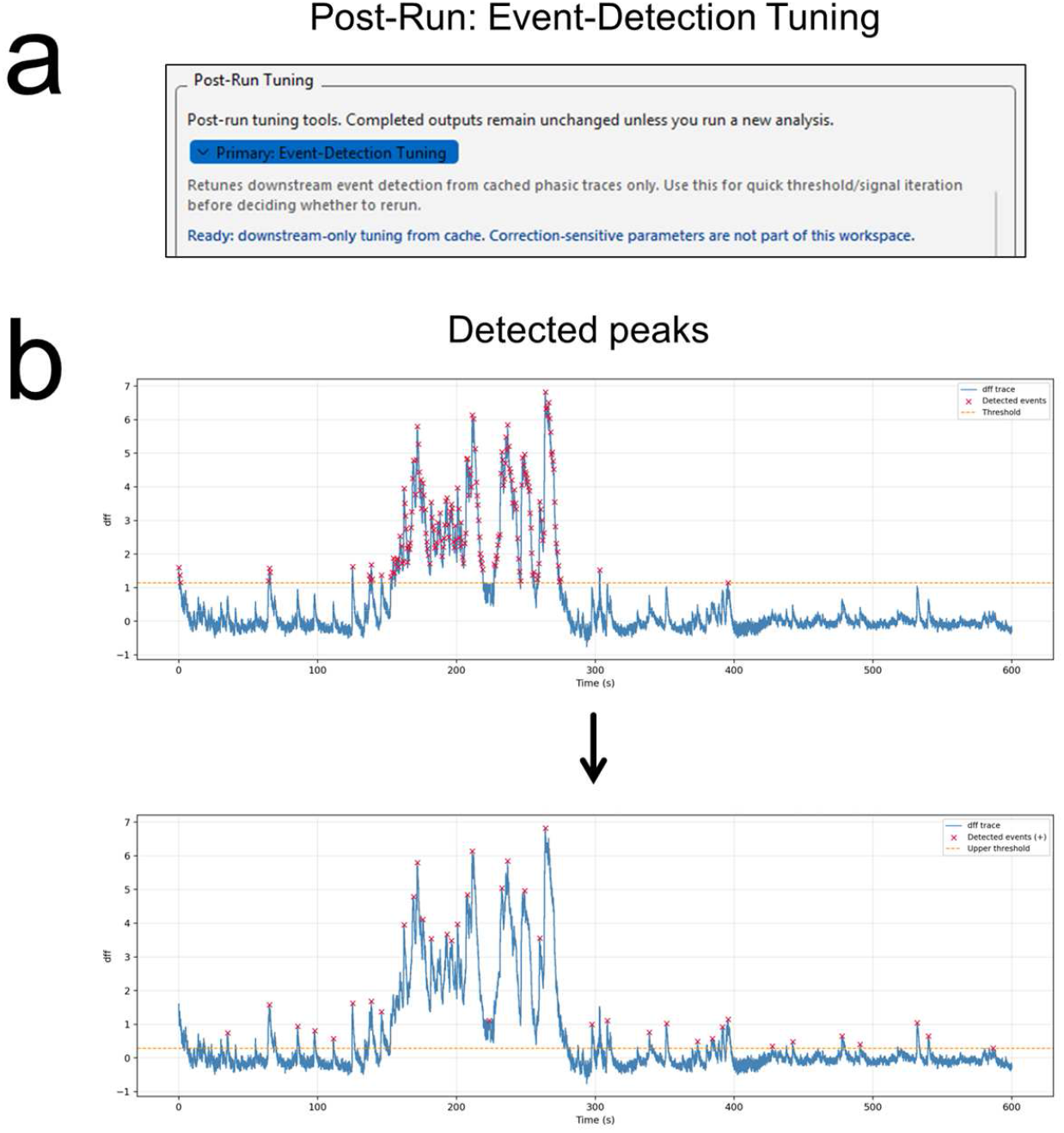
Post-run retuning of downstream event analysis. **a)** Post-run tuning window for downstream event-analysis retuning, performed from cached phasic traces after completion of the initial run. **b)** Representative phasic ΔF/F trace before and after detection retuning. In the top panel, the initial settings produce excessive peak detection. In the bottom panel, the adjusted settings yield a more appropriate set of detected events from the same trace.

### Multiscale visualization of long-term outputs

The same run can be inspected across multiple timescales and outputs in paired views (**Fig. 6**). In the representative example, centered signal and isosbestic traces, dynamic fit output, and corrected ΔF/F are each displayed in a format that preserves multiday temporal organization while also allowing closer inspection of individual recording intervals. This makes it possible to examine raw channel structure, correction behavior, and corrected output within the same run. Multiday organization and local signal structure can therefore be examined together.

**Figure 6.**
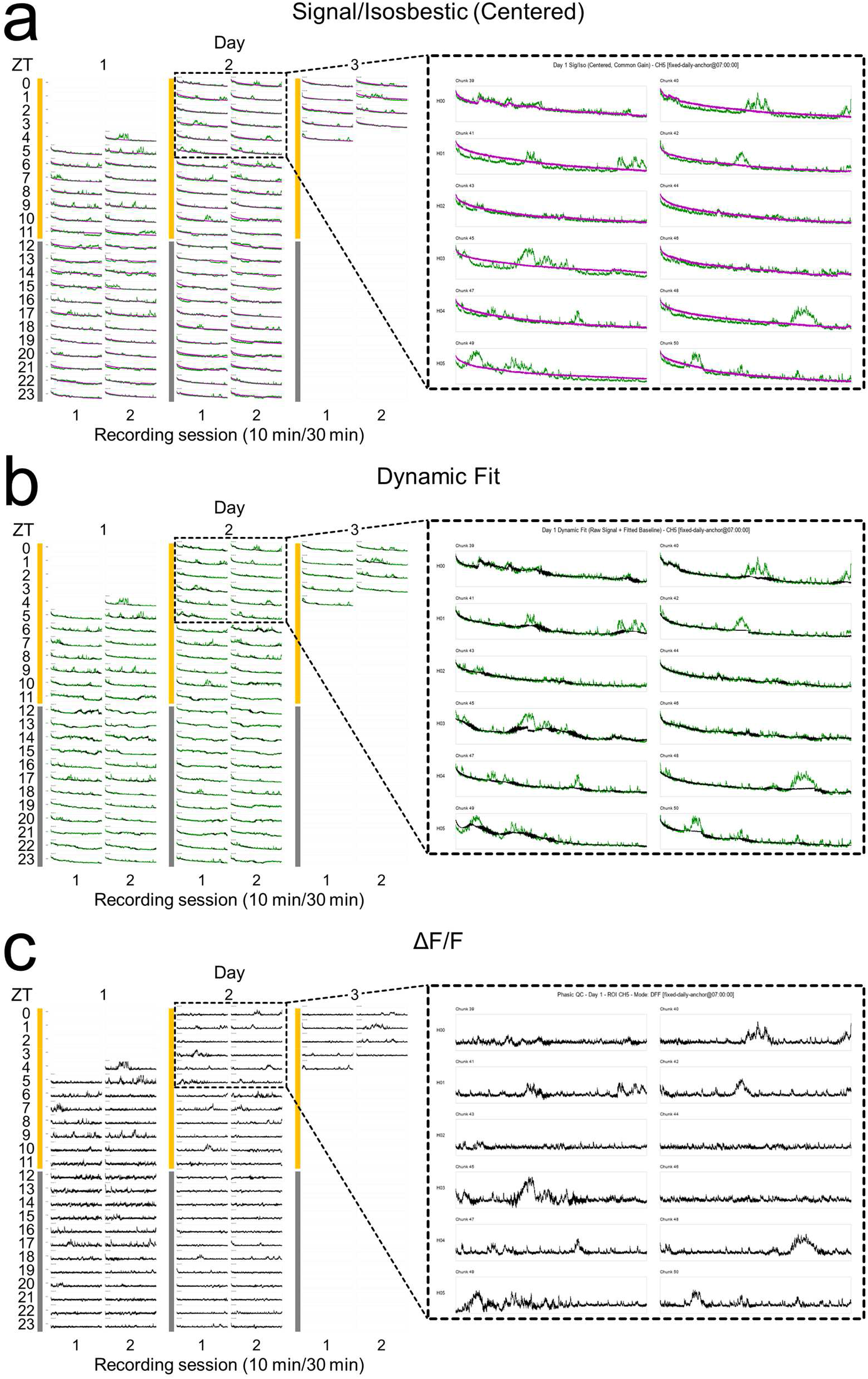
Stacked and expanded views of analysis outputs. **a-c)** Day-aligned stacked displays of a 48 h recording arranged on a three-day zeitgeber time grid, with two 10 min sessions per hour. Because recording began at ZT 4.5, early slots on Day 1 and late slots on Day 3 are intentionally blank, preserving alignment to the 24 h structure. Dashed boxes and connectors indicate the interval expanded at right. **a)** Centered signal and isosbestic traces. **b)** Dynamic fit output. **c)** Corrected ΔF/F.

### Complementary tonic and phasic outputs for long-term analysis

Tonic and phasic outputs capture different features of long-term recordings (**Figs. 7**,**8**). Tonic outputs preserve slow baseline structure that would not be captured by phasic event summaries alone. In these tonic output views, recordings with relatively weak and strong slow oscillatory structure can be distinguished over the course of the recording (**Fig. 7**).

**Figure 7.**
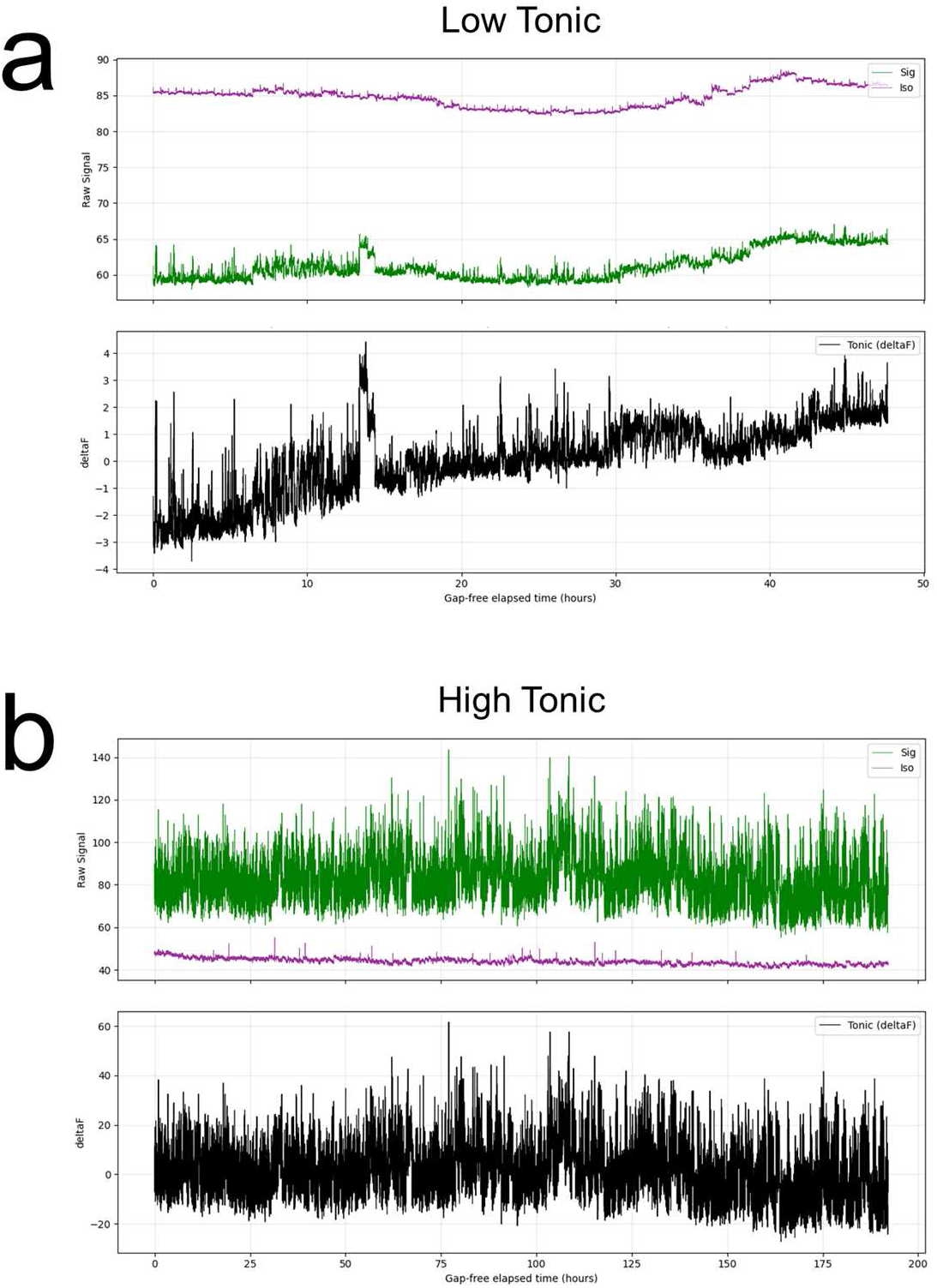
Tonic outputs from low- and high-tonic recordings. **a)** Tonic output window from an animal with a low tonic oscillation, shown as signal and isosbestic traces over elapsed recording time (*top*) and the corresponding ΔF trace (*bottom*). **b)** Tonic output window from an animal with a high tonic oscillation, shown in the same format. For visualization, session-level bleaching was flattened, session baselines were preserved, and intermittent recording sessions were plotted gap-free across elapsed time.

Phasic outputs provide complementary summaries of event occurrence and signal magnitude over time (**Fig. 8**). Event frequency reports how often detected events occur, whereas integrated fluorescence reports how much above-threshold signal accumulates over the same time axis. These summaries distinguish event rate from cumulative signal magnitude and complement the slower baseline structure captured by tonic outputs.

**Figure 8.**
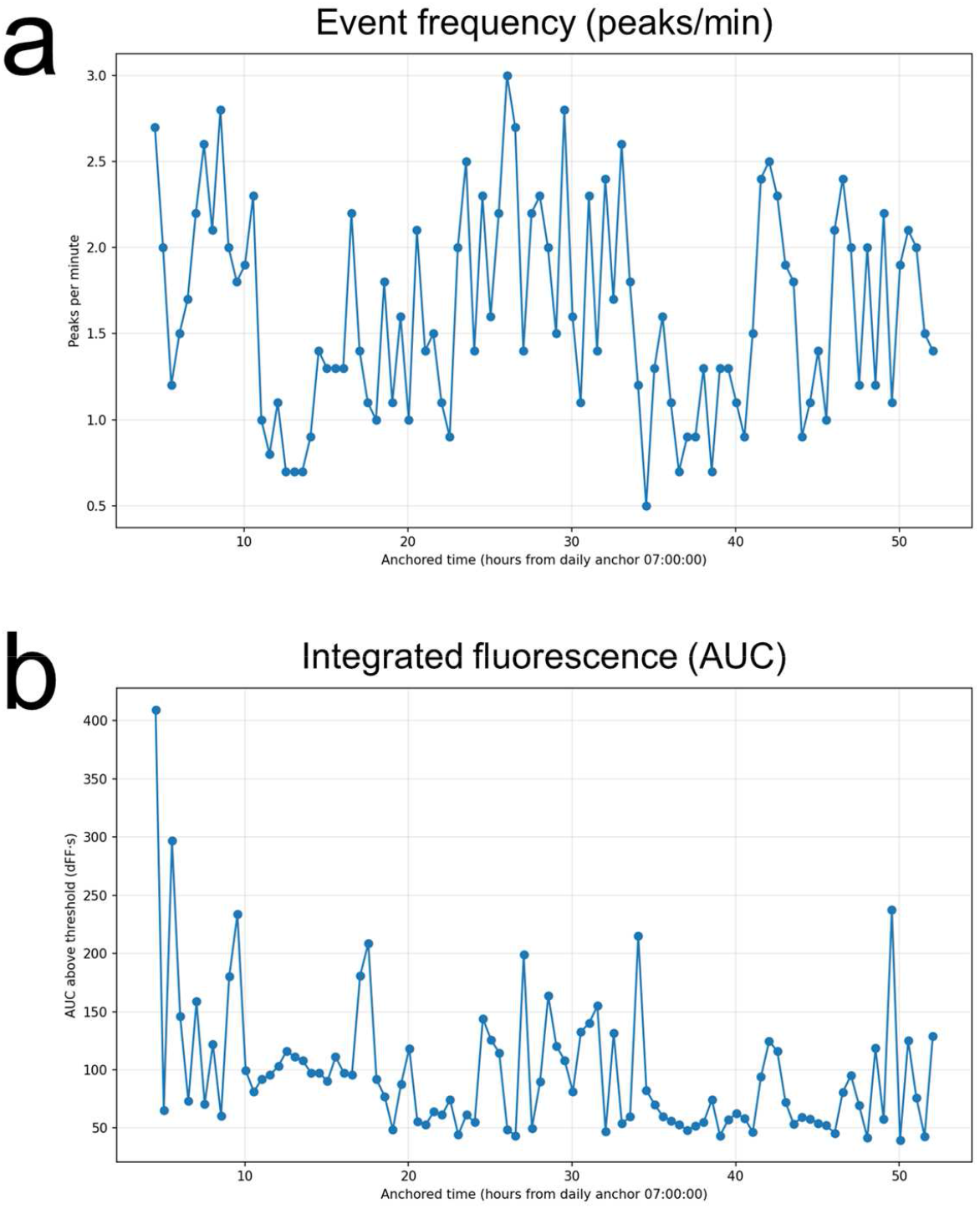
Event frequency and integrated fluorescence over time. **a)** Event frequency plotted as peaks per minute across anchored time, with the x axis aligned to the daily anchor at 7:00 AM (ZT 0). **b)** Integrated fluorescence plotted across the same anchored time axis, quantified as area under the curve above threshold (ΔF/F·s).

## DISCUSSION

Long-term fiber photometry requires an analysis workflow built for recordings that extend across hours to days or weeks [1]. The present study provides a practical software environment for long-term fiber photometry analysis in which runs are configured, validated, executed, inspected, and then reopened through distinct post-run branches for correction, retuning and downstream event reanalysis. The software also preserves multiple analytical views of the same recording, including tonic outputs, correction-level inspection, corrected phasic outputs, and downstream phasic summaries, while allowing the same run to be inspected at both multi-day and session-level scales.

This software is best understood in the context of existing open-source photometry analysis toolkits. Packages such as GuPPy, pMAT, and PyFiber have lowered the barrier to entry for photometry analysis, particularly for short-term experiments and event-triggered workflows [16–18]. However, these methods were developed around several assumptions including short recordings, single-session file structures, and stimulus-locked analyses. Those assumptions become limiting for long-term datasets where meaningful information may be present in both slow tonic variation and in phasic activity over time. Long-term analysis therefore requires scalable handling of large session collections, preservation of tonic structure that might otherwise be removed as baseline drift, robust and flexible isosbestic fitting, and outputs that remain interpretable across multiple timescales. The present software implements those requirements in a workflow designed for photometry recordings spanning hours to days.

In that context, several aspects of the software are especially important. The examples shown here demonstrate that isosbestic correction choice can substantially change the corrected signal itself. Correction is therefore treated as a revisitable analytical decision. Downstream event-analysis settings can likewise be revised independently from upstream correction. The software also preserves tonic and phasic outputs as distinct but related views of the same recording, which matters because slow baseline structure may be biologically meaningful and phasic summaries alone provide only a partial account of signal dynamics [20]. Visualization is also part of the analytical workflow: representative signal and isosbestic traces, broad multiday views, and stacked session-aligned outputs help determine whether a recording is dominated by tonic structure, phasic activity, or a mixture of both [9,21]. In our software, these outputs are available within the same environment used for run setup, execution, and post-run revision, making it easier to inspect preprocessing choices, evaluate correction behavior, and interpret local waveform structure in the context of the broader temporal pattern of the recording.

The present software also has important limits. We do not claim that the correction modes shown here are universally optimal across all sensors, circuits, or recording conditions, or that this workflow is equally appropriate for every photometry application. Instead, we aim to provide a reproducible and revisitable environment in which correction choices, event detection settings, and related analysis settings can be made deliberately and inspected directly. In doing so, the software addresses the central requirements for long-term photometry analysis, including scalability, revisitable correction, distinct tonic and phasic outputs, and transparent multiscale inspection. Those capabilities will be essential if the field is to move beyond lab-specific custom pipelines toward a more generalizable and trusted analytical methodology.

## METHODS

### Animals and housing

Male wild-type (n = 1) and ChAT-IRES-Cre (n = 1) mice, 3 months old, on a C57BL/6J background contributed the recordings used in this study [22]. Mice were singly housed at constant temperature (∼23°C) and humidity (∼40%) under either a 12 h:12 h light:dark cycle (LD; lights on defined as zeitgeber time 0) or in constant darkness (DD), with food and water available ad libitum. All procedures were approved by the Texas A&M University Institutional Animal Care and Use Committee and performed in accordance with institutional guidelines.

### Stereotaxic surgery

Wild-type mice received unilateral hippocampal injections of AAVDJ/8-hSyn-GCaMP7s (∼4.40 × 10^12^ GC/mL; University of Zurich Viral Vector Facility). ChAT-IRES-Cre mice received unilateral basal forebrain injections of AAVDJ/8-hSyn-DIO-GCaMP7s (∼8.80 × 10^12^ GC/mL; University of Zurich Viral Vector Facility) [23]. Mice were anesthetized with isoflurane, and buprenorphine-SR (1 mg/kg, s.c.) was administered before surgery. After the skull was leveled, AAVDJ/8-hSyn-GCaMP7s was injected into the hippocampus (-2.00 mm posterior, +1.25 mm lateral, -1.25 mm ventral from bregma) at a volume of 100 nL and a rate of 1 nL/min. In ChAT-IRES-Cre mice, AAVDJ/8-hSyn-DIO-GCaMP7s was injected into the basal forebrain (+0.85 mm anterior, 0.00 mm lateral, -4.35 mm ventral from bregma) at a volume of 100 nL and a rate of 1 nL/min. The injection needle was left in place for 10 min before withdrawal. A fiber optic cannula (2 or 6 mm length, 200 μm core diameter, 0.39 NA; RWD Life Science) was implanted at the corresponding injection site and secured with adhesive cement (Parkell) and opaque black dental cement. The implant was covered with a thin layer of black nail polish. Mice recovered in their home cages for 2 weeks before experiments.

### Fiber photometry

Fiber photometry data were collected using a multi-channel fiber photometry system (R821; RWD Life Science). After postsurgical recovery, mice were tethered to the photometry system and allowed at least 2 d to acclimate to the patch cable (200 μm core diameter, 0.37 NA, 3 m length; Doric Lenses) before recordings began. Fluorescence was sampled at 40 frames/s using interleaved 470 nm (signal) and 415 nm (isosbestic) excitation channels with a 33% duty cycle (10 min on, 20 min off).

### Long-term recording structure

Representative examples in this study were drawn from multiday photometry recordings acquired under the duty cycle described above. Depending on the output shown, recordings were visualized either in elapsed time or after alignment to zeitgeber time or a daily anchor. Outputs included tonic views, corrected phasic signals, stacked multiday displays, and anchored phasic summaries.

### Software analysis workflow

All analyses shown here were performed using the Python-based software environment presented in this study. Runs were configured and validated within the GUI before tonic and phasic analyses were executed. Completed runs were then inspected through the Analysis Output interface, and representative examples of correction retuning and downstream event reanalysis were produced using the corresponding post-run tuning workflows.

## Code and software availability

Code and software are available at https://github.com/jones-lab-tamu/long-term-photometry.

## ACKNOWLEDGMENTS

This work was supported by National Institutes of Health Grant R35GM151020 (J.R.J.), a research grant from the Whitehall Foundation (J.R.J.), and National Institutes of Health Grant F31DA065512 (J.N.C.). We thank members of the Jones Lab for helpful discussions and feedback on the manuscript.

## REFERENCES

1. Tang Q, Cook JN, Yurgel ME, Hattar S, Jones JR. Long-term optical monitoring of genetically-encoded fluorescent indicators. PNAS Nexus. 2025;4: pgaf372.

2. Vas S, Wall E, Zhou Z, Kalmar L, Han SY, Herbison AE. Long-term recordings of arcuate nucleus kisspeptin neurons across the mouse estrous cycle. Endocrinology. 2024;165: bqae009.

3. Hrvatin S, Sun S, Wilcox OF, Yao H, Lavin-Peter AJ, Cicconet M, et al. Neurons that regulate mouse torpor. Nature. 2020;583: 115–121.

4. Vandendoren M, Landen JG, Rogers JF, Killmer S, Alimiri B, Pohlman C, et al. Oxytocin neurons signal state-dependent transitions from rest to thermogenesis and behavioral arousal in social and non-social settings. Neuroscience. bioRxiv; 2024. Available: https://www.biorxiv.org/content/10.1101/2024.10.22.619715v3.full.pdf

5. Rinker JA, Hoffman M, Knapp J, Wukitsch TJ, Kutlu MG, Calipari ES, et al. Prelimbic neuron calcium activity predicts perceived hedonic value across drinking solutions and ethanol dependent states in mice. bioRxiv. 2023. p. 2023.04.04.535635. doi:10.1101/2023.04.04.535635

6. Yaguchi K, Hagihara M, Konno A, Hirai H, Yukinaga H, Miyamichi K. Dynamic modulation of pulsatile activities of oxytocin neurons in lactating wild-type mice. PLoS One. 2023;18: e0285589.

7. Chen F, Dong X, Wang Z, Wu T, Wei L, Li Y, et al. Regulation of specific abnormal calcium signals in the hippocampal CA1 and primary cortex M1 alleviates the progression of temporal lobe epilepsy. Neural Regen Res. 2024;19: 425–433.

8. Douglass AM, Kucukdereli H, Madara JC, Wang D, Wu C, Lowenstein ED, et al. Acute and circadian feedforward regulation of agouti-related peptide hunger neurons. Cell Metab. 2024. doi:10.1016/j.cmet.2024.11.009

9. Jones JR, Simon T, Lones L, Herzog ED. SCN VIP Neurons Are Essential for Normal Light-Mediated Resetting of the Circadian System. J Neurosci. 2018;38: 7986–7995.

10. Jones JR, Chaturvedi S, Granados-Fuentes D, Herzog ED. Circadian neurons in the paraventricular nucleus entrain and sustain daily rhythms in glucocorticoids. Nat Commun. 2021;12: 5763.

11. Gomez AM, Wu Y, Zhang C, Boyd L, Wee T-L, Gewolb J, et al. Aberrant hypothalamic neuronal activity blunts glucocorticoid diurnal rhythms in murine breast cancer. Neuron. 2025;0. doi:10.1016/j.neuron.2025.11.019

12. Sayar-Atasoy N, Aklan I, Yavuz Y, Laule C, Kim H, Rysted J, et al. AgRP neurons encode circadian feeding time. Nat Neurosci. 2023. doi:10.1038/s41593-023-01482-6

13. Yurgel ME, Gao C, O’Malley JJ, Tang Q, Yanay N, Bashford AR, et al. A stress-sensing circuit signals to the central pacemaker to reprogram circadian rhythms. Sci Adv. 2025;11: eadr7960.

14. Simpson EH, Akam T, Patriarchi T, Blanco-Pozo M, Burgeno LM, Mohebi A, et al. Lights, fiber, action! A primer on in vivo fiber photometry. Neuron. 2023. doi:10.1016/j.neuron.2023.11.016

15. Enoki R, Ono D, Kuroda S, Honma S, Honma K-I. Dual origins of the intracellular circadian calcium rhythm in the suprachiasmatic nucleus. Sci Rep. 2017;7: 41733.

16. Sherathiya VN, Schaid MD, Seiler JL, Lopez GC, Lerner T. GuPPy, a Python toolbox for the analysis of fiber photometry data. bioRxiv. 2021. p. 2021.07.15.452555. doi:10.1101/2021.07.15.452555

17. Bruno CA, O’Brien C, Bryant S, Mejaes J, Pizzano C, Estrin DJ, et al. pMAT: An Open-Source, Modular Software Suite for the Analysis of Fiber Photometry Calcium Imaging. bioRxiv. 2020. p. 2020.08.23.263673. doi:10.1101/2020.08.23.263673

18. Conlisk D, Ceau M, Fiancette J-F, Winke N, Darmagnac E, Herry C, et al. Integrating operant behavior and fiber photometry with the open-source python library Pyfiber. Sci Rep. 2023;13: 16562.

19. Keevers LJ, Jean-Richard-Dit-Bressel P. Obtaining artifact-corrected signals in fiber photometry via isosbestic signals, robust regression, and dF/F calculations. Neurophotonics. 2025;12: 025003.

20. Zhai Q, Zeng Y, Gu Y, Li Z, Zhang T, Yuan B, et al. Time-restricted feeding entrains long-term behavioral changes through the IGF2-KCC2 pathway. iScience. 2022;25: 104267.

21. Tang Q, Godschall E, Brennan CD, Zhang Q, Abraham-Fan R-J, Williams SP, et al. Leptin receptor neurons in the dorsomedial hypothalamus input to the circadian feeding network. Science Advances. 2023;9: eadh9570.

22. Rossi J, Balthasar N, Olson D, Scott M, Berglund E, Lee CE, et al. Melanocortin-4 receptors expressed by cholinergic neurons regulate energy balance and glucose homeostasis. Cell Metab. 2011;13: 195–204.

23. Zhang Y, Rózsa M, Liang Y, Bushey D, Wei Z, Zheng J, et al. Fast and sensitive GCaMP calcium indicators for imaging neural populations. Nature. 2023;615: 884–891.

